# Male-male interactions select for conspicuous male colouration in damselflies

**DOI:** 10.1101/2020.06.24.168823

**Authors:** Md Kawsar Khan, Marie E. Herberstein

**Affiliations:** Department of Biological Sciences, Macquarie University, Sydney, Australia

**Keywords:** Ontogenetic colour change, Sexual selection, Sensory ecology, Communication and signalling, Colour polymorphism, Sexual conflict, Scramble competition

## Abstract

Male ornamentation, such as conspicuous male colouration, can evolve through female mate choice. Alternatively, in species without overt female mate preference, conspicuous male colouration can evolve via intrasexual selection to resolve male-male competition or to prevent costly male-male mating attempts. Here, we investigated the drivers of conspicuous male colouration in an ontogenetic colour changing damselfly, *Xanthagrion erythroneurum*, where the juvenile males are yellow and change colour to red upon sexual maturity. We first showed that red males were chromatically and achromatically more conspicuous than the yellow males. We then quantified the condition of the males and showed that red males were larger and in better condition than yellow males. We tested female preference in a choice experiment where we artificially manipulated male colour, and found that females did not choose mates based on male colouration. We further tested whether the male colouration affected male-male interactions. We presented red and yellow males in the breeding arena, and found that red males received less intra- and interspecific aggression than yellow males. Our study experimentally showed, for the first time, that male conspicuousness is not a target of female mate choice in damselflies. Intra- and interspecific male-male interactions therefore appear to be the driver of conspicuous male colouration in damselflies.

## Introduction

Male armaments and ornaments, such as horns in antelopes, the long tails of peacocks, giant hind legs in beetles, and conspicuous colours in birds, lizards and insects, are not thought to afford survival benefits, but rather evolve through sexual selection (Balmford et al., 1992a; Balmford et al., 1992b; Emlen, 2008; Parker, 2013; Petrie et al., 1991). Among ornamental traits, the function and evolution of male ornamental colouration has been studied in many taxa since Darwin proposed his sexual selection theory (Jones and Ratterman, 2009). Male conspicuous colouration can evolve via intersexual selection through female preference (Gomez et al., 2009; Kemp, 2007), or via intra-sexual selection through male-male competition for mating and/or to avoid male-male mating attempts (Bajer, Molnár, Török, & Herczeg, 2011; Sherratt, 2001).

Females are predicted to express mate choice if males vary in their ability to provide either direct benefits such as parental care, nuptial gifts, higher social status, safer territory, or indirect benefits such as higher immunity and better genetic alleles (Albo and Peretti, 2015; Georgiev et al., 2015; Kirkpatrick and Barton, 1997; López Pilar and Martín José, 2005). In this context, conspicuous colouration may function as a signal of male condition (Hill, 1991; Keyser and Hill, 2000) and by selecting to mate with a conspicuous males over dull males females may be responding to signals of good condition (Montoya and Torres, 2015; Setchell, 2005; Vásquez and Pfennig, 2007). Consequently, conspicuous male colour can evolve via female mate choice.

Conspicuous male colouration can evolve via male-male competition, in addition to or without female preference, when males compete for limited breeding resources such as nest sites, oviposition sites, or to access females for mating (Ahnesjö et al., 2001; Klug et al., 2010; Morris et al., 1992; Wacker and Amundsen, 2014). Competition for breeding resources can largely determine the mating success in some species, therefore sexual selection can favour male traits to acquire breeding resources (Debuse et al., 2003; Shuster and Wade, 2003). Otherwise, male-male competition to access females arises when the males outnumber females, or when the frequency of receptive females in the breeding territory is lower than that of males (Klug et al., 2010; Weir et al., 2011). Because of a male biased sex ratio, the males compete with each other to access females, and only the best quality males are assumed to secure mates (Forslund, 2000; Morris et al., 1992). Conspicuous colouration can function as an honest signal of male condition, resource holding potentiality and fighting ability (Ligon and McGraw, 2013; Lim and Li, 2013; Weaver et al., 2017). Conspicuous males may therefore maintain larger and safer breeding territories, higher social dominance, and better access to females, which result in higher mating success (Korzan and Fernald, 2007; Setchell and Wickings, 2005).

Conspicuous male colouration is common in many damselflies (Corbet 1999; Bybee et al. 2016). In territorial damselflies, conspicuousness is condition-dependent and related to males maintaining territory and achieving higher mating success via female mate choice and inter- and intraspecific male-male competition (Siva-Jothy, 1999; Svensson et al., 2004; Tynkkynen et al., 2005). The function of conspicuous male colouration is, however, largely debated in damselflies where the males do not defend territory. In these damselflies, adult male and female damselflies assemble in waterbodies such as ponds, lake, streams for mating, and ovipositing. Because of the high male density and limited oviposition sites, males compete to access and persist in the mating territories. In such scramble scenarios, males approach females from behind for mating and females cannot see the colour of males, in the first step of the mating sequence, which is unlikely to be the point of mate selection (Corbet 1999; Sherratt and Forbes 2001). Subsequently, however, males require the cooperation of females to lock genitalia (Fincke, 1997) and it has therefore been argued that conspicuous male colour can evolve through female mate choice.

Male-male competition for mating is the other probable function of conspicuous male colouration in damselflies with scramble mating system. In these damselflies, a large number of conspecific and heterospecific males gather within a breeding arena (pond) for mating, resulting in male biased aggregations (Corbet, 1999). Conspecific male-male competition arises when two or more males compete over access to the same female or breeding spot. On the other hand, interspecific male-male competition occurs in sympatric populations to acquire limited oviposition and mating sites. These intra- and interspecies competition could induce male-male interactions which can reduce male fitness (Gering, 2017). In such circumstances, conspicuous male colouration can evolve to signal male condition thereby reducing intra- and interspecific male-male interactions. This hypothesis, although intriguing, is yet to be experimentally tested.

Here we tested the causative agents of the conspicuous male colouration in *Xanthagrion erythroneurum* damselflies. *Xanthagrion erythroneurum* exhibits ontogenetic colour change, where the males change colour from yellow to red, about a week after their emergence (Khan and Herberstein, 2020). First, we tested if red males are more conspicuous than yellow males for damselfly vision. We then assessed whether the conspicuous colour if conspicuous colouration signals male condition in this species. Finally, we experimentally tested whether female mate choice or male-male interactions select for the conspicuous red colouration in males. We predicted the females would prefer red males over yellow males if the conspicuous red colouration evolves through female mate choice. On the other hand, male-male interactions select for conspicuous male colouration we predicted that red males would receive less aggression less conspecific and heterospecific aggression than yellow males in the breeding arena.

## Methods and Materials

### Study Species

*Xanthagrion erythoneurum* is a medium-sized damselfly (21-23mm) belonging to the Coenagrionidae family (Zygoptera: Odonata). This species is widely distributed throughout Australia and commonly found in stagnant freshwater reservoirs such as ponds, marshes, lakes and dams. The adult males can be distinguished from other sympatric damselflies by their red face, red thorax, the red colouration of the first and second abdominal segments and the blue bands on the eighth and ninth abdominal segments (Khan and Herberstein, 2019; Theischinger and Hawking, 2016). In the Sydney region, this species starts emerging in September and is seen in flight until June (Khan and Herberstein, 2020). During this whole period, this species remains reproductively active (Khan and Herberstein, 2020).

### Field site

We collected the damselflies from and carried out experiments at a pond located on the North Ryde campus of Macquarie University, Sydney, Australia (33.772 S, 151.114 E). In this pond, *Xanthagrion erythroneurum* cooccurs with other damselflies including *Ischnura heterosticta, Austroagrion watsoni, Austrolestes annulosus, Diplacodes melanopsis* and *Orthetrum caledonicum*. The sympatric species *Ischnura heterosticta* and *Austroagrion watsoni*, like *Xanthagrion erythroneurum*, do not defend a territory but searches for females in the ponds which looks more like a scramble competition for mating. All three species share shoreline vegetations at ponds for perching and mating and submerged vegetations for ovipositing.

### Reflective spectrometry

We measured the reflective spectra of the collected males and leaves of the vegetation surrounding the pond to quantify the visual background using a JAZ EL-200 portable spectrophotometer (Ocean Optics, USA) with a PX-2 pulsed light source. We measured all spectra in a dark room by setting the spectrophotometer to a constant boxcar width and integration time settings of 10 and 20 milliseconds respectively, and to average five scans. We measured the reflectance relative to a white standard (Ocean Optics, USA) to standardize the measurements. We first immobilized the damselflies by placing them in a refrigerator at 4°C for five minutes. Then, we set the probe of the spectrophotometer perpendicular to the cuticular surface of the metathorax from a fixed working distance of two millimetres. We took three reflectance spectra of each male and each background leaf from 300 nm to 700 nm and subsequently averaged those three measurements. We processed the reflectance spectra with OceanOptics Spectrasuite software (ver. 1.6.0_11) and eventually binned to one nm wavelength intervals before minor LOESS smoothing (α = 0.35). We performed the spectral processing using the package ‘pavo’ v 2.1 (Maia et al., 2019) in R v 3.5.2 (R core team 2018).

### Visual modelling

We calculated the chromatic and achromatic contrast of the red and yellow males against their background using the receptor noise model (Vorobyev and Osorio, 1998; Vorobyev et al., 2001). This model calculates the detectability between two colours in just noticeable difference (JND) units where one JND means that the receiver can distinguish between the colours (Vorobyev et al., 2001). The receptor noise model has previously been applied in behavioural studies to predict colour discriminability in various taxa including odonates (Barry et al., 2015; Huang et al., 2014; Khan and Herberstein, 2019).

We aimed to determine how the colour and luminescence of the red and yellow males are perceived by the receiver i.e. conspecific and heterospecific damselflies. The visual system of *X. erythroneurm* is not known. We however, know that damselflies of the Coenagrionidae family can have either trichromatic or tertachromatic visual system (Henze et al., 2013; Huang et al., 2014). We therefore applied both systems to calculate the chromatic and achromatic contrast of the red and yellow males against their backgrounds (Khan and Herberstein, 2019). For trichromatic visual modelling, we applied the photoreceptor sensitivities of *Ischnura heterosticta*, while the photoreceptor sensitivities of *Ischnura elegans* were used for tetrachromatic modelling (Henze et al., 2013; Huang et al., 2014). We used the photoreceptor sensitivities of these two species as they are the closest related species in the phylogenetic tree of our study system whose visual system is known.

We calculated the quantum catches of the photoreceptors by following the methods of Vorobyev and Osorio (1998) (Supplementary method S1). We used standard daylight (D65) as the ambient light spectrum. We then log-transformed the quantum catches according to the Weber-Fencher law (Vorobyev et al., 2001). Finally, we calculated the chromatic and achromatic contrast of red and yellow males against their background as a function of the log-transformed quantum catches weighted by the noise of each photoreceptor (Vorobyev and Osorio, 1998). We performed the visual modelling in R v 3.5.2 (R core team, 2018) using the package pavo v 2.1 (Maia et al., 2019).

### Male condition

We calculated body length, body mass, lipid and protein content to determine the condition of the males (Castaños et al., 2017). We captured *X. erythroneurum* males from the field using an insect sweep net (Khan, 2015; Khan, 2018) and brought them back to the Behavioural Ecology Laboratory at Macquarie University for morphometric measurements. We took measurements of the damselflies within two hours after collecting them. We weighed the body mass of the live damselflies on a balance (Mettler toledo, accuracy 0. 01 mg). Next, we immobilized the damselflies by cooling them in a refrigerator at 4°C for five minutes. We then positioned the damselflies laterally and took digital photographs using a Canon 600D camera mounted with Canon EF 55-250 lens. We measured the total body length of the damselflies from the digital photographs using the ImageJ software (Schneider et al., 2012).

### Lipid Quantification

We measured the lipid content of the damselflies by the gravimetric method (Barry and Wilder, 2013). First, we euthanised the damselflies by placing them in a −30°C freezer for 10 minutes, then we dried the damselflies at 60°C for 48 hours and then weighed their dried body mass. We then submerged the dried damselflies in chloroform. After 24 hours, we discarded the chloroform, and replaced it with fresh chloroform for another 24 hours. The chloroform was then discarded and the damselflies were air-dried under a fume hood at room temperature for another 24 hours. We further dried the damselflies for another 24 hours at 60°C and later reweighed the dried damselflies. The lipid content of each damselfly was calculated as the difference between the body mass of the damselfly before and after chloroform extraction.

### Protein extraction and quantification

We extracted the soluble protein from the damselflies using 0.1 M NaOH (Sigma-Aldrich) as a lysis buffer. First, we finely ground the dried damselflies with a polypropylene pestle and added 0.1 N NaOH (100μl per 1mg of insect weight). We then vortexed the lysate, sonicated it for 30 minutes in a water bath, and then heated at 90°C for 15 minutes. Finally, we centrifuged the lysate at 13000 rpm for 10 minutes, discarded the undigested tissues that precipitated, and collected the clarified lysate from the supernatant.

We quantified the protein content in the clarified lysates of the damselflies using Pierce™ BCA protein assay kit (Thermo Fisher Scientific). We used the Bovine Serum Albumin (BSA) supplied with the protein assay kit for preparing the standard solution. We used a linear range of standard BSA protein concentrations from 1.35 mg to 0.05 mg for making the standard absorbance curve. We used 0.1 M NaOH (Sigma-Aldrich) as a diluent for the standard solution preparation, for the damselfly lysates preparation and also for the blank control. We took 25 μL of standard BSA for standard solution preparation and 25 μL damselfly lysates for sample protein quantification and then added 0.1 M NaOH (Sigma-Aldrich) to make the final volume 200 μL. We added the standard and sample solution in triplicates in a in 96 well flat-bottomed plate and incubated at 37 °C for 30 min. We took the absorbance of the incubated plates at 562 nm using a FLUOstar OPTIMA microplate reader (BMG Labtech). We quantified the relative protein quantity of the lysates using the standard absorbance curve.

### Female mate choice experiment

We experimentally tested if females prefer mating with red or yellow males. The yellow males are sexually immature and unable to mate (Khan and Herberstein, 2019). We, therefore, painted red males with yellow colour to determine the effect of male yellow colour on female mate choice. We performed the female mate choice trials by restraining a female in an insect mating cage (58cm × 32cm × 34cm) with a natural red male and a red male that was painted yellow. We manipulated the thorax of the damselflies by painting yellow over red using non-toxic Tim & Tess™ poster paint (Khan and Herberstein, 2019). We also painted red over the natural red to control for the effect of paintings. We took spectra of the painted damselflies to approximate their natural colour (supplementary figure S1).

We placed the mating cage approximately three meters away from the pond — the natural habitat of the damselflies. We recorded the sexual interactions of the damselflies while sitting one meter away from the cages. We counted the number of tandems formed by the control red males and the manipulated yellow males. The tandem is the first step of damselfly mating where a male becomes physically connected to a female by his cerci. A tandem event can dissociate if the female does not cooperate, or it can form a wheel if the female cooperates. When a tandem was formed, we continued the trials until the tandem disassociated or formed a wheel. We recorded the duration of tandems when disassociated and when forming a wheel. When a wheel was formed, we recorded the duration of the wheel before disassociation.

We performed the female mate choice trials on sunny days between 10:00 hrs and 16:00 hrs when mating usually occurs in the field (Khan and Herberstein, 2019). We performed each trial for 30 minutes with two new males and a female; no damselflies were reused. Paint was washed off the damselflies after every trial, and the damselflies were released at the end of the day. The aim of the experiment was to determine female mate choice between the red and yellow males, so in the analysis we included trials where a female choose to form a tandem or wheel with one of the males.

### Male-male interactions experiment

We conducted three sets of experiments to determine the effect of male colour on male-male interactions by tethering the experimental males. In the first experiment, we tethered a naturally occurring red male and a naturally occurring yellow male and determined the male-male interactions received by the red and yellow males. To determine whether the incurred interactions are the effect of colour or other developmental changes, we painted a red male yellow and tethered it with a red male and determined any male-male interactions. In the third experiment, we altered the colour of yellow males by painting them red and presented the naturally occurring yellow males with the red-painted yellow males and determined any male-male interactions. If body colour determines the interactions at the pond, we expect the yellow painted red males will receive less aggression than the red males. Similarly, the red-painted yellow males will receive less aggression than yellow males.

We applied a modified damsel-on-a-dowel technique (Fincke, Fargevieille, & Schultz, 2007) to determine the male-male interaction of the red and yellow males in their natural habitat. We glued a live yellow male and a red male 20 cm apart from each other on a dowel using UHU^™^ glue. The damselflies were glued in perching positions, with their legs attached to the dowel. The dowel was then placed at the edge of the pond. In this pond, *X. erythoneurum* coexists with two heterospecific damselfly species of the Coenagrionidae family: *Ishnura heterosticta and Austroagrion watsoni*, with whom they share mating and oviposition sites. We measured the aggressive and non-aggressive responses received by the red and yellow *X. erythoroneurum* males from conspecific males and heterospecific (*Ishnura heterosticta*) males. We observed the responses by sitting approximately one meter away from the dowel, which allowed us to observe the focal damselflies clearly without disturbing regular movements of the approaching damselflies. An approaching damselfly can detect the focal damselfly when it passes within 10 cm of the focal damselfly (Fincke, 2015) and can either show aggression or non-aggression when it passes. When an intruder male passed within 10 cm left or right of the focal male without any physical contact, we counted it as a non-aggressive interaction. On the other hand, when the intruder male bit the experimental male, we counted it as an aggressive interaction (Fincke et al., 2007). Finally, if the intruder male tried to form a clasp (grab the focal male and tried to move its cerci to the prothorax of the focal male) or formed a tandem (intruder male physically connected with the focal male), we counted it as a mating attempt.

We conducted each trial for 10 minutes by placing the tethered damselflies in different locations around the lake. We used the same pair for three consecutive trials unless the focal damselflies were predated by sympatric odonates after one or two trials. We conducted all our experiments on sunny days between 10:00 hrs and 1600 hrs when damselfly density and interactions are high (Khan and Herberstein, 2019). We counted aggressive and non-aggressive responses received by the focal males from approaching conspecific and heterospecific males. We manipulated damselfly colour following the same procedure as described in the female mate choice experiment. After finishing the experiment, we unglued the damselflies by hand, washed off the paint, and released them at the end of the day.

### Statistical analyses

We applied Shapiro-Wilk tests to determine normality and F-tests to compare the variance of the data. Two-sample t-tests were applied to compare the chromatic and achromatic contrast of the males against their background. We applied Two-sample t-tests to analyse the total length, and Welch Two Sample t-test to analyse body mass, protein content and lipid content of red males and yellow males – Bonferroni corrections were applied to adjust the p-values.

We applied Generalized linear models (GLMs) to determine whether females are more likely to form tandems and wheels with red males than yellow painted red males. We fitted GLMs with the numbers of tandems and wheels as a response variable and male colour (red or painted yellow) as covariates. We applied Cox regression models to analyse tandem duration and wheel duration of the red and yellow painted red males. Tandems that transitioned to wheels were analysed separately from tandems that dissociated before forming wheels.

To analyse the aggression received by red and yellow males in male-male interaction experiments, we applied generalized linear mixed models (GLMMs) with aggression or non-aggression as a response variable, focal male (red male and yellow) and intruder male (conspecific male and heterospecific male) as fixed effects and the pair identity as a random effect. For the experiments involving natural red and yellow males and natural red males with a red male painted yellow we applied binomial distribution. On the other hand, we applied a quasi-binomial distribution (to account for the over-dispersion) for the experiment in which a natural yellow male with a yellow male painted red were used as focal male. For each analysis, we used the full model (glmer (attack, passby) ~ focal male*intruder male + (1|id)) by including interactions between fixed effects and pair identity as random effect. All the analyses were conducted in R v 3.5.2 (R core team, 2018) using the ‘survival’ (Therneau and Lumley, 2019), ‘lme4’ (Bates et al., 2019), and ‘MuMIn’ (Bartoń, 2019) packages.

## Results

### Damselfly spectra

The reflectance spectra of the yellow males showed peaks between 588 nm and 700 nm whereas the red males showed peaks between 657 nm and 700 nm (Figure 1a). The reflectance spectra of the background showed a Gaussian peak between 551-554 nm (Figure 1a).

**Figure 1:**
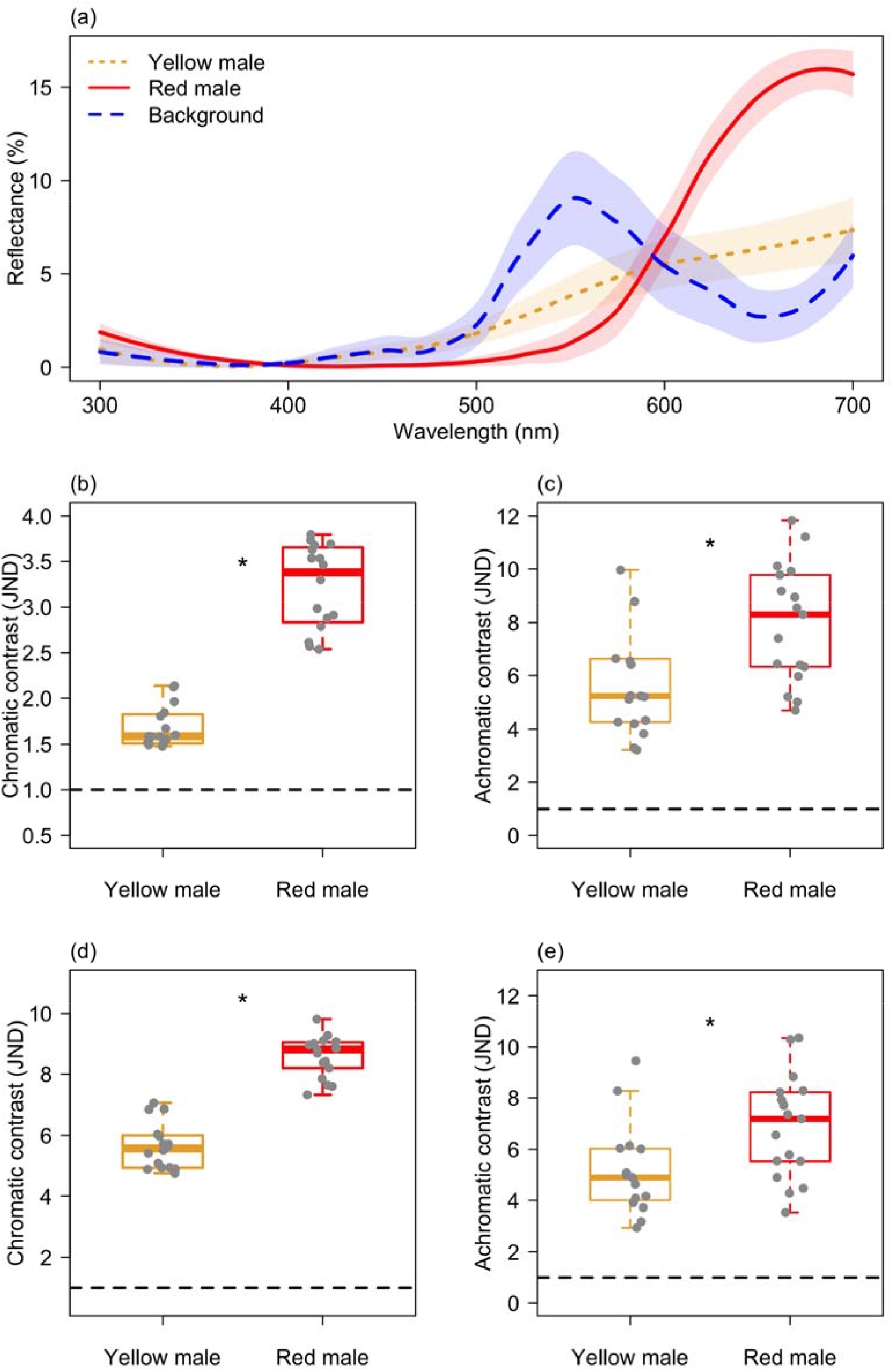
a) Reflectance spectra (mean ± standard deviation) of the red males (*n* = 17), yellow males (*n* = 17) and background leaves (*n* = 30); b) chromatic and c) achromatic contrast of the red and yellow males against their background in trichromatic damselfly vision; d) chromatic and e) achromatic contrast of the red and yellow males in tetrachromatic damselfly vision. Boxplots depict the median, the 25th and 75th percentile with the whiskers extended to the minimum and maximum data points. Outliers that were > 1.5 times the interquartile range. (* indicates p <0.05).

### Visual modelling

The chromatic and achromatic contrast between the red and yellow males and the background was more than one JND, suggesting that the males are discriminable against their background by both trichromatic (Figure 1b–1c) and tetrachromatic (Figure 1d–1e) visual systems (Figure 1b–1e). The red males showed both higher chromatic contrast (Mann-Whitney U test: *W* = 0, *p* <0.0001) and achromatic contrast (Welch two sample t-test: *t* = −2.07, *df* = 27.32, *p* <0.05) than the yellow males in trichromatic damselfly vision (Figure 1b–1c). Similarly, in tetrachromatic damselfly vision, the red males showed higher chromatic contrast (Two sample t-test: *t* = −11.81, *df* = 31, *p* <0.0001) and achromatic contrast (Two sample t-test: *t* = −2.49, *df* = 30, *p* <0.05) than the yellow males (Figure 1d–1e).

### Male condition

The red males were longer in total length (Two sample t-test: *t* = −5.13, *df* = 75, *p* <0.0001) and their body mass was heavier (Welch two sample t-test: *t* = −16.65, *df* = 70.39, *p* <0.0001) than the yellow males (Figure 2a–2b). Furthermore, lipid content of the red males was significantly higher (Mann-Whitney U test: *W* = 21, *p* <0.0001) than in yellow males (Figure 2c). Similarly, protein content of the red males was significantly higher (Welch two sample t-test: *t* = −8.69, *df* = 28.25, *p* <0.0001) than in yellow males (Figure 2d).

**Figure 2:**
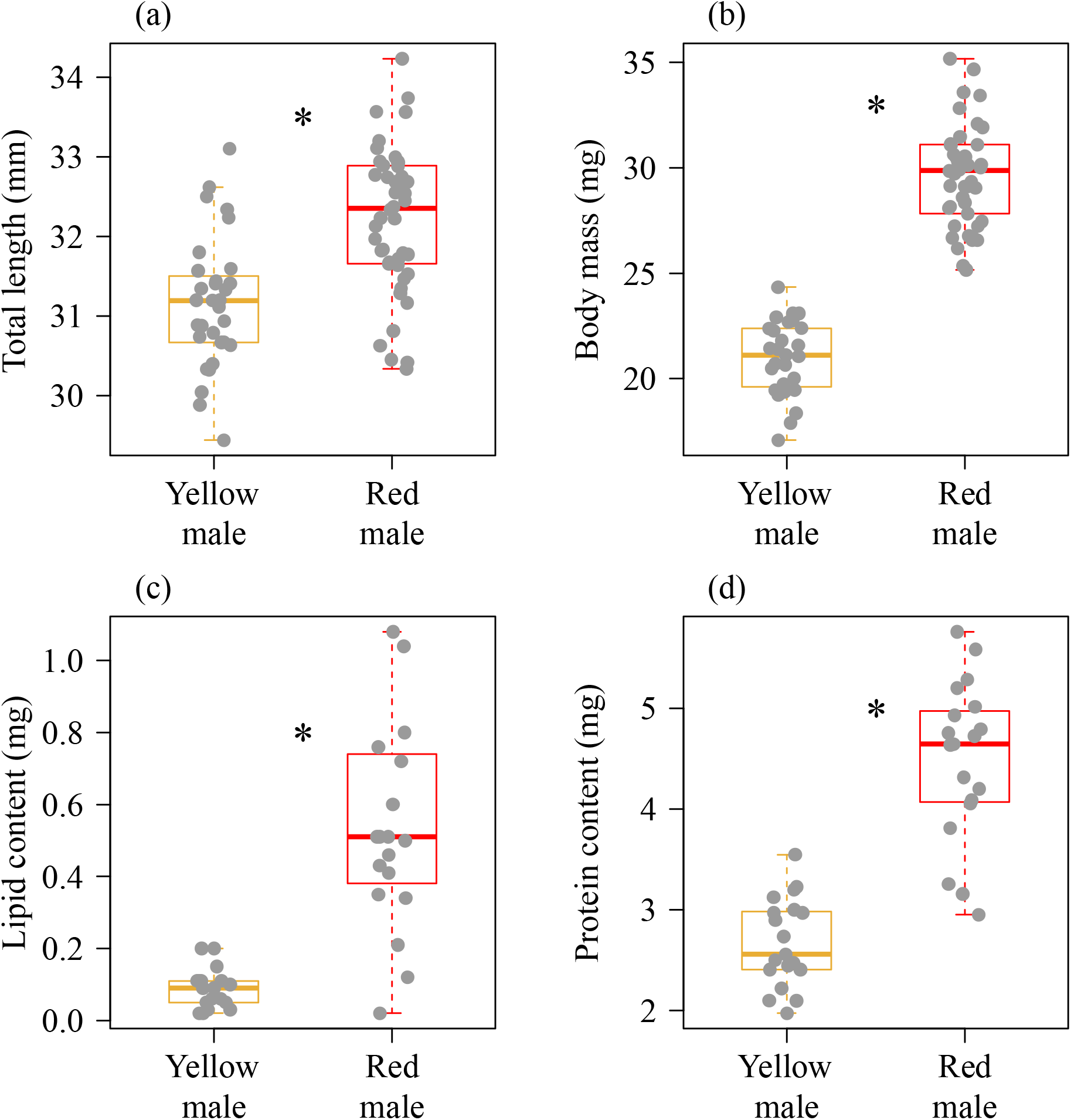
a) Total length of the yellow males (*n* = 31); b) and red males (*n* = 46); b) body mass of the yellow males (*n* = 27) and red males (*n* = 46); c) lipid content of the yellow males (*n* = 19), and red males (*n* = 19), and d) protein content of the red males (*n* = 19) and yellow males (*n* = 19). Boxplots depict the median, the 25th and 75th percentile with the whiskers extended to the minimum and maximum data points. Outliers that were > 1.5 times the interquartile range were excluded. (* indicates p <0.0001).

### Female mate choice

In total, 60 tandems were formed during the female mate choice experiment. The number of tandem formations did not differ significantly (GLM: *χ^2^* = 0.46, *df* = 1, *p* = 0.49) between the red males (*n* = 32) and the yellow painted red males (*n* = 28) (Figure 3a). Over half (51.7%) of the males failed to form a wheel after forming a tandem. The number of males proceeding from tandem to wheel did not differ significantly (GLM: *χ^2^* = 1.63, *df* = 1, *p* = 0.20) between the red males (*n* = 13) and the yellow painted red males (*n* = 16) (Figure 3b). When the tandems dissociated before forming wheels, the tandem duration was not affected by male colour (Cox regression: *χ^2^* = 1, *df* = 1, *p* = 1) (Figure 3c). Similarly, the time to wheel formation did not differ significantly (Cox regression: *χ^2^* = 0.62, *df* = 1, *p* = 0.6) between the red or yellow painted red males (Figure 3d). Finally, the duration of the wheel before disassociation did not differ significantly (Cox regression: *χ^2^* = 0.06, *df=* 1, *p* = 0.06) between the red males (*n* = 13) and the yellow painted red males (*n* = 16) (Figure 3e).

**Figure 3:**
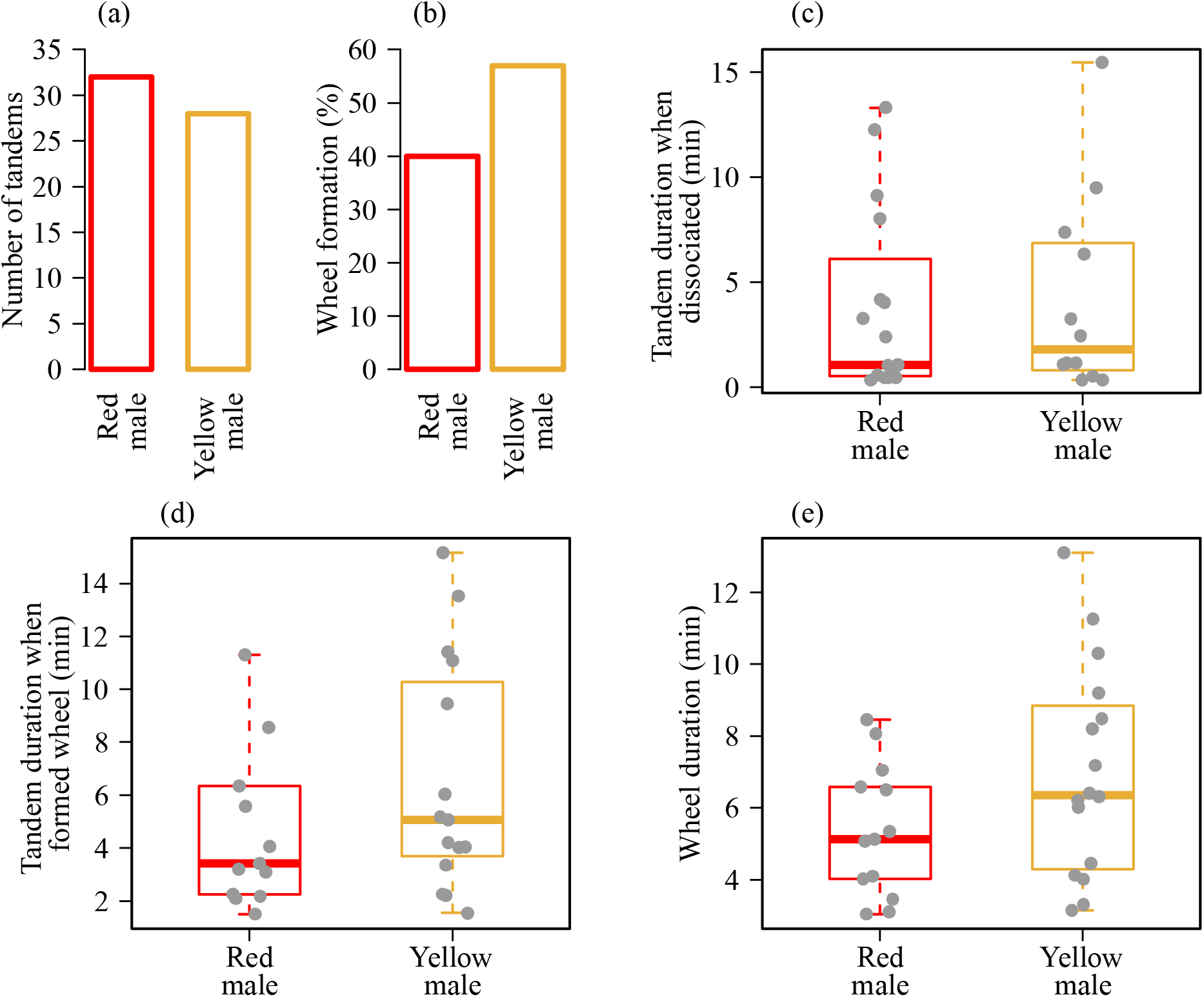
a) Number of tandems formed by red males (*n* = 32), and yellow males (*n* = 28); b) percentage of the tandems involving red males (*n* = 13) and yellow males (*n* = 16) that ended in wheel formation; c) duration of red and yellow male tandems that ended in dissociation rather than wheel formation; d) duration of red and yellow male tandems that ended in wheel formation, and e) wheel duration of red and yellow males Boxplots depict the median, the 25th and 75th percentile with the whiskers extended to the minimum and maximum data points. Outliers that were > 1.5 times the interquartile range were excluded.

### Male-male interactions

The yellow males received significantly higher aggressive responses than red males from conspecific males (estimate: 4.48 ± 0.38, z = 11.74, p < 0.001) in the male-male competition experiment when natural red and yellow males were presented to intruder males (Figure 4a). Also, there was a significant interaction between focal males and intruder males (estimate: −1.59 ± 0.67, *z* = −2.37, *p* < 0.05) showing that the probability of receiving aggression from heterospecific males was higher in yellow males than red males (Figure 4b; supplementary table 2). Similarly, when yellow painted red males were presented with natural red males, yellow painted red males received significantly higher aggression than natural red males from conspecific and heterospecific males (Figure 4c-d; supplementary Table 3-4). Furthermore, when yellow males were painted red, and presented with the natural yellow males, the natural yellow males received higher aggression from conspecific and heterospecific males than the red painted yellow males (Figure 4e-f; supplementary table 5-6).

**Figure 4:**
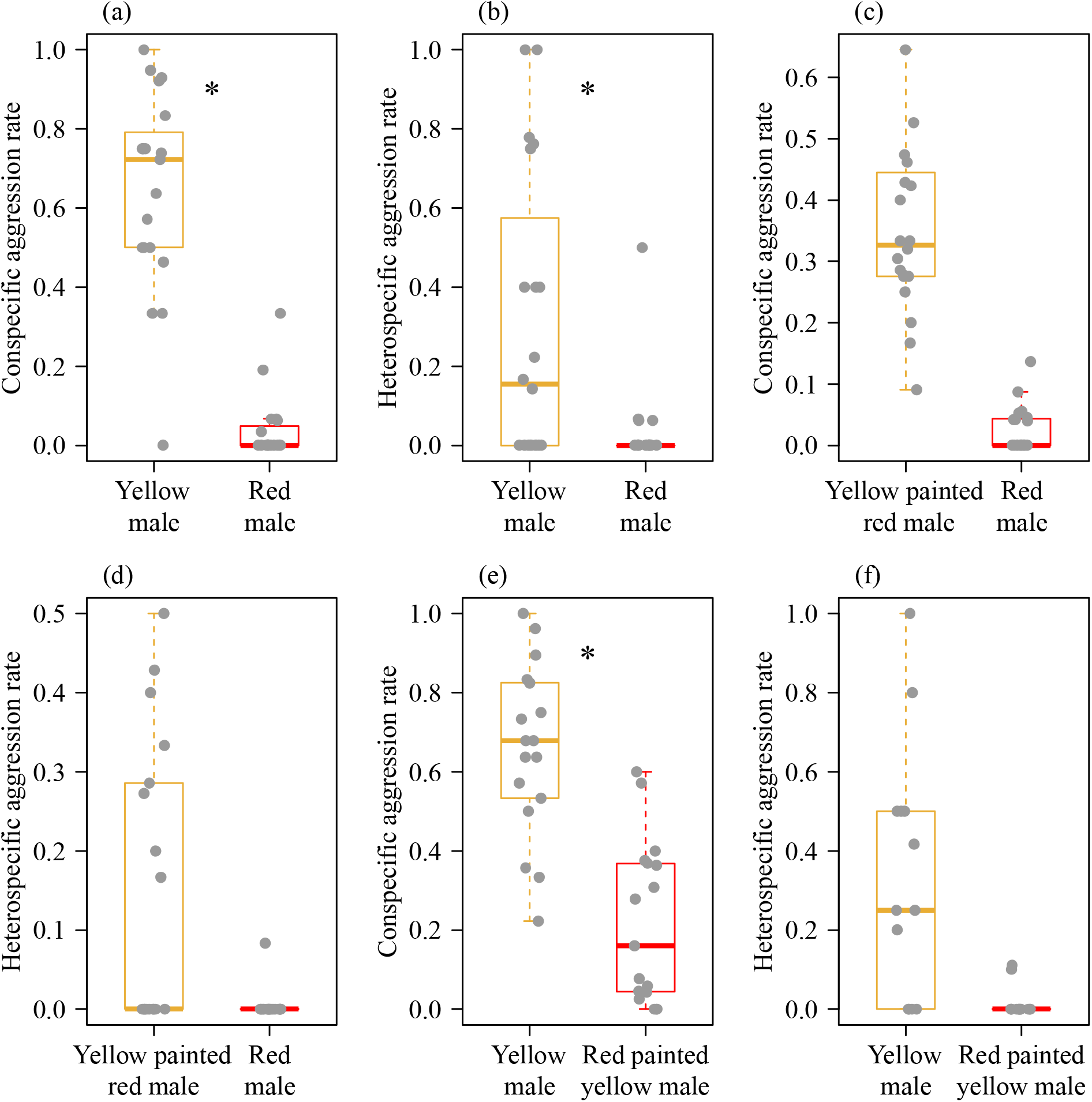
Aggressions (number of attacks/number of approaches) received by natural red and natural yellow males (*n* = 20) from a) conspecific males and b) heterospecific males; c) aggressions received by natural red males, and yellow painted red males (*n* = 20) from conspecific males and d) heterospecific males; e) aggressions received by natural yellow males and red painted yellow males (*n* = 18) from conspecific males, and f) heterospecific males. Boxplots depict the median, the 25th and 75th percentile with the whiskers extended to the minimum and maximum data points. Dots are data points; outliers that were > 1.5 times the interquartile range were excluded.

## Discussion

Conspicuous male colouration can evolve through female mate choice, male-male competition for mating, to reduce male-male mating attempts or through a combination of all three (Clutton-Brock, 2007; Sherratt & Forbes, 2001). We investigated the drivers of conspicuous male colouration in *Xanthagrion erythroneurum* damselflies. We found that red males were chromatically and achromatically more consipicuous than yellow males. We further showed that red males were in better nutritional and physiological condition than yellow males. Next, we experimentally tested female preferences for male colouration and found that the females did not prefer red males over yellow males when given a choice. Finally, we tested male-male interactions in the breeding arena and found that yellow males received more aggression than red males from conspecific and heterospecific males.

The female mate choice experiments showed that the number of tandems did not differ between red and yellow males. In the cage experiment, females cannot avoid tandem as they cannot fly away from the approaching males. A female, however, can reject a mate by dissociating from the tandem (Khan and Herberstein, 2019). In support of that, we found that the females rejected 51.7% of mating attempts in our trials. The rejection rate, however, was not different between the red and the yellow males. A female can further express refusal by delaying the wheel formation, or by dissociating from the wheel quickly before sperm transfer (Khan and Herberstein, 2019). If females preferred red males, we would have expected 1) tandem durations before dissociation are longer for red males, 2) red males attain wheel more quickly, and 3) the wheel durations are longer for red males. We, however, did not find significant differences between the red and the yellow males in any of these choice indicators. Females are unlikely to detect the coloration of the males as they approach from behind to form a tandem. The females probably use tactile cues and clasping strength rather than colour to estimate male quality (Barnard and Masly, 2018; Barnard et al., 2017). Taken together, our results strongly suggest that female preferences are not selective agents of male colouration. Cryptic female choice, however, could select for male conspicuous colouration, and further studies are required for test this possibly.

Conspicuous male colouration can evolve by intrasexual selection if conspicuousness increases male mating success by reducing conspecific aggression in the breeding arena. Thus, we predicted that red males would receive less aggression than yellow males. Our results confirmed this prediction: yellow males (whether natural or painted) received more overall aggression than painted or natural red males. The red males received less aggression probably because the red colour signals male quality and competitive ability, thereby, serving as a status badge to resolve costly disputes without direct physical contact. Our findings are consistent with previous findings suggesting that red colouration is a signal of male condition and dominance, and functions to intimidate rivals in lizards (Healey et al., 2007; Whiting et al., 2006), fishes (Dijkstra et al., 2005), birds (Pryke and Griffith, 2006) and primates (Setchell and Wickings, 2005). Furthermore, we showed that red painted yellow males received more aggression than control yellow males supporting the hypothesis that red is inherently intimidating to rivals (Baird et al., 2013; Barlow, 1983; Pryke, 2009; Rowland et al., 1995) even when additional phenotypic information (e.g. size) was available.

Interspecific interactions can be a significant evolutionary force to shape traits in sympatric species (Tynkkynen et al., 2004; Tynkkynen et al., 2005). Here, we showed that the natural and painted red males received less heterospecific aggression than the natural and painted yellow males. The heterospecific aggression can occur due to an interspecific recognition error where males are phenotypically similar or because of male competition for common resources. Our study species, Xanthagrion erythroneurum with a red thorax and *Ischnura heterosticta* with a blue thorax are phenotypically dissimilar, therefore recognition error is probably an unlikely mechanism for the observed interspecific aggression. On the other hand, both species assemble at the pond for breeding, share the same perching sites for mating, foraging, and resting in between mate searching, and also share the same oviposition sites suggesting interspecific competition for breeding resources are a possible mechanism. Our findings support the idea that conspicuous colouration can reduce interspecific aggression (Drury and Grether, 2014) to acquire shared breeding resources (Lipshutz, 2018; Peiman and Robinson, 2010). In breeding ponds where damselflies mate, a large number of conspecific and heterospecific males aggregate and compete for limited mating and oviposition sites (Corbet, 1999). This suggests that, in *X. erythroneurum*, and probably also in other damselflies, the conspicuous colouration has evolved to reduce intra- and interspecific male-male interactions.

Conspicuous male colouration can also evolve through intrasexual selection to avoid costly male-male tandem attempts (Sherratt, 2001). This anti-harassment aposematic hypothesis has been supported in moor frogs showing that the males attain conspicuous colouration upon reproductive maturity to avoid male-male amplexus formation (Sztatecsny et al., 2012). Similarly, sexually dimorphic abdominal blue bands in non-territorial damselflies reduce intraspecific male-male tandem formation (Beatty et al., 2015; Khan and Herberstein, 2019). In the male-male competition experiment of this study, while presenting the males, we found that conspecific males formed tandems with 11 out of 20 experimental yellow males, but never with the red males (data not shown). This preliminary result suggests that the conspicuous red colouration might also function to avoid male-male tandem formation in this species. Further behavioural choice experiments to test conspecific male mate choice between the red and yellow morphs are needed to better understand the anti-harassment signal of the red colour.

Our study combined colour vision modelling with laboratory and behavioural experiments to explain the function of the conspicuous colouration in damselflies with a scramble mating system. We showed that the conspicuous red colour in *X. erythroneurum* damselflies signal male condition. We further demonstrated that the conspicuous colouration is not a target of female mate choice but reduces inter- and intraspecific male aggression in the breeding arena. Our findings suggest that conspicuous male colouration can evolve tto reduce costly male-male interactions. Because our study presents clear intra-sexual advantage of red males, the question of why yellow males persist in the population remains elusive. Further studies are needed to explain if being yellow is a resource limited constrain or it is an adaptation to reduce the foraging and predation risks associated with a red colour.

## Acknowledgements

We thank Jim McLean for his comments on the initial version of the manuscripts. MKK thanks Payal Barua for all her support. MKK designed the study, collected and analysed the data, and wrote the manuscript. MEH contributed to the design of the study, and critical revision of the manuscript.

## Data accessibility

All data will be uploaded in Figshare upon acceptance for publication

